# stTrace: Detecting Spatial-Temporal Domains from spatial transcriptome to Trace Developmental Path

**DOI:** 10.1101/2025.05.19.654812

**Authors:** Zhangdi Song, Changyu Zheng, Jiaxing Chen

**Affiliations:** Guangdong Provincial/Zhuhai Key Laboratory IRADS, Beijing Normal-Hong Kong Baptist University, 2000 Jintong Road, Tangjiawan, Zhuhai, 519087, Guangdong Province, China; Hong Kong Baptist University, Kowloon Tong Kowloon, 100084, Hong Kong, China

**Keywords:** Spatial-temporal domain, Development, Trajectory, spatial transcriptome

## Abstract

Development is essential for the growth and functional maintenance of organisms. Investigating the development process is vital for uncovering the formation of complex biological systems. However, current approaches to studying development from gene expression rely primarily on single-cell gene expression data to infer developmental trajectories, neglecting the spatial distribution of cells within tissues and their interactions. Although spatial transcriptomics provides spatial context for gene expression, existing algorithms focus mainly on identifying spatial regions without further exploring their developmental connections. In this study, we propose an algorithm for detecting spatial-temporal domains in tissue to trace developmental path (stTrace) using spatial transcriptomics. stTrace integrates the degree of cell development, gene expression, and spatial location to identify “spatial-temporal domains”, regions where cells share similar functions and developmental stages within the tissue. Moreover, hierarchical relationships exist among these regions, reflecting developmental connections between cells in the tissue. Our experiments on mouse embryo and human breast cancer data revealed that stTrace can detect more refined regions than traditional spatial domain identification algorithms. Furthermore, the directions of developmental paths inferred from hierarchical relationships are consistent with the dynamic trajectories derived from single-cell velocity.

**Key points:**

- We proposed spatial-temporal domain, which refers to spatially continuous regions where cells exhibit similar gene expression patterns, related functions, and close developmental levels.
- We present stTrace, a computational framework for identifying spatial-temporal domains by iteratively optimizing two objectives: minimizing Structure Entropy to capture functional similarity and hierarchy, and maximizing the Silhouette Coefficient to ensure temporal compactness.
- In both mouse embryo and human breast cancer datasets, spatial-temporal domains align better with H&E-stained images and and have better compactness in developmental level.
- stTrace detected developmental paths that are well-mapped with cell’s temporal information like single-cell velocity and trajectories and application show the usage in studying embryo development and cancer invasion.

## Introduction

Understanding how organisms form and differentiate is essential for advancing developmental biology and disease pathology [3, 26, 8, 13]. In multicellular organisms, processes such as cell replication, differentiation, and development occur under spatial constraints [32, 17, 16, 19, 14]. Additionally, gene expression within cells follows a distinct spatial and temporal pattern [20, 25, 1], ensuring that processes such as cell growth, division, and differentiation occur coordinated. Therefore, exploring the relationship between cell developmental level, gene expression, and spatial location is essential for understanding organogenesis, disease progression, and regenerative medicine.

To study development, Spateo [23] constructs morphometric vector field by comparing the gene expression profiles of cells at corresponding locations across different time points, predicting the spatial migration patterns of single cells and tracking the morphological changes of individual organs over time. However, morphometric vector field analysis requires gene expression profiles from multiple time points and only emphasizes the cell migration direction without further discussing the overall hierarchical structural organization. Another category approach to studying developmental processes mainly relies on trajectories analysis [35]. Researchers have proposed various methods for inferring the trajectory from single cell. For example, Monocle [31] use independent component analysis and a minimum spanning tree to construct differentiation trajectories, while Monocle2 [22] employs principal component analysis and reversed graph embedding. Wolf et al. [36] introduced Partition-based graph abstraction, which uses dimensionality reduction and topology to generate network sketches and analyze connections between cell subpopulations. These methods assume that similarity in gene expression profiles reflects the developmental level of cells, without considering the spatial constrain and the interactions with neighboring cells. Although stLearn [21] has introduced a pseudo-space-time trajectory analysis algorithm, which attempted to integrate spatial information into cell trajectory inference. However, stLearn only incorporates Euclidean distance in a weighted manner into the overall distance calculation, which does not discuss the integration of the two types of information thoroughly and, therefore is still limited. In this work, we propose to better integrate spatial information with gene expression to analyze development and reveal the hierarchical structural organization in tissue.

On considering gene expression and spatial information, researchers have proposed methods for detecting spatial domains from spatial transcriptomics [28], where spatial domain refers to spatially continuous regions in which cells exhibit similar gene expression patterns or related biological functions. For example, MULTILAYER [18] applies clustering in continuous locally defined transcriptomics and adopted community detection methods for graphical partitioning; DeepST [37] uses a deep learning framework to identify spatial domains; BayesSpace [40] enhances the resolution with a Bayesian statistical approach; STAGATE [7] learns low-dimensional latent embedding with integrating the information of spatial and gene expression. Even though there are various spatial domain detection methods nowadays, these methods would be insufficient for studying hierarchical structural organization that implies the development process, due to the lack of consideration of the developmental level and neglecting potential links between domains.

To better incorporate spatial information and gene expression patterns into inferring the development, we propose a novel algorithm, stTrace, for spatial-temporal domain detection and development path reconstruction. The spatial-temporal domain refers to spatially continuous regions where cells exhibit similar gene expression patterns, related functions, and close developmental levels (Fig. 1). To measure cell signature from the temporal aspect, we calculated the Signaling Entropy Rate for each cell to reflect the development level. On the spatial and functional aspects, stTrace generates the region partition based on the enhanced gene expression profile’s similarity with optimizing the Structure Entropy. Here, Structure Entropy is selected for spatial partition assessment because it shows how well the hierarchical relationship between partitions is, which reflects the hierarchy organization along the development process. On the other hand, the Silhouette Coefficient assesses the consistency and separability of the cells in the temporal dimension. Subsequently, the two types of signals are integrated by consensus from bootstrap experiments. Then iterative refinement is applied to improve border as well as region partition in temporal, spatial and functional aspects further, facilitating the identification of spatial-temporal domains. Finally, stTrace reconstructs the developmental path based on the hierarchical relationships among these spatial-temporal domains.

**Fig. 1.**
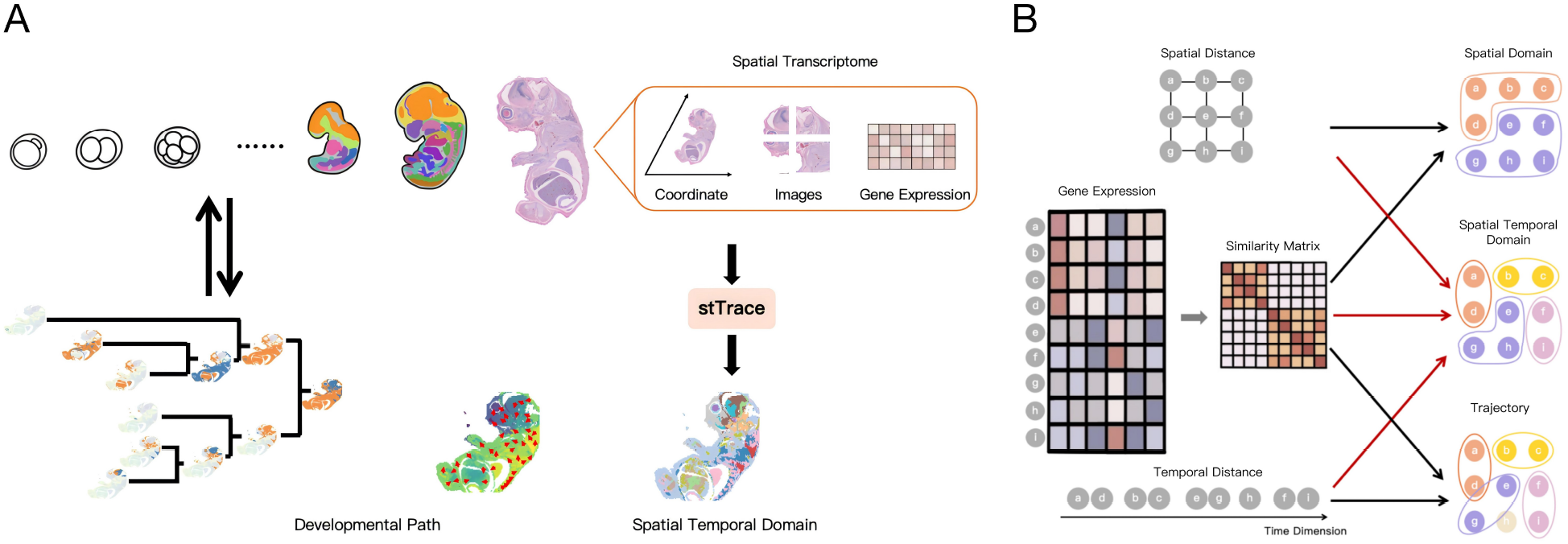
(A). The graphic abstract of stTrace. We take the typical one-time point slice when organisms have gone through development. Spatial transcriptome gets coordinate, image, and gene expression information from the slice. Based on the above information, stTrace can reveal the spatial-temporal domain and infer the development path that maps to the history of organism development. (B). A comparison of spatial domain, spatial-temporal domain, and trajectory. Spatial domain methods use gene expression data and spatial coordinates (x, y) to delineate static tissue regions, capturing spatial continuity but ignoring temporal progression; Trajectory inference methods reconstruct developmental paths based solely on gene expression and pseudotime distances, effectively modeling cellular progression over time but neglecting spatial context; Spatial-temporal domain methods integrate gene expression, spatial location, and developmental information to identify domains that are coherent across both space and time dimension.

We applied stTrace to study the development of mouse embryos and breast cancer from Visium Spatial Gene Expression. The result shows that stTrace can detect domains with clearer regional boundaries that better align with the staining patterns of samples, and n higher temporal compactness with larger inter-difference and smaller intra-difference, compared to existing spatial domain identification algorithms. Moreover, the reconstructed developmental path in mouse embryo exhibits consistency with developmental stages implied by Signaling Entropy Rate [29, 30], Pseudotime [22], Single Cell Velocity [2] and MOSTA [5], which is a previous study on mouse development from spatial transcriptome on multiple time slices. In the human breast cancer experiment, the developmental path identified by stTrace shows the invasive direction of cancer cells, and the signals are also verified by the distribution of cancer marker genes. In addition, the ablation study demonstrates that incorporating developmental information enables effective identification of cells that are spatially adjacent and functionally similar but at distinct developmental level. In conclusion, stTrace offers a novel approach to study development by effectively combining spatial, functional and temporal aspects.

## Methodology

stTrace aims to identify the spatial-temporal domain and reconstruct the developmental path using the development degree inferred from spatial transcriptome data, simultaneously constraining on spatial and functional similarity. In the beginning, we generated an enhanced expression matrix from the spatial transcriptome data, which includes gene expression profiles, spatial coordinates, and corresponding H&E-stained images (As shown in Fig. 2A, details about data preprocessing are described in Supplementary Materials).

**Fig. 2.**
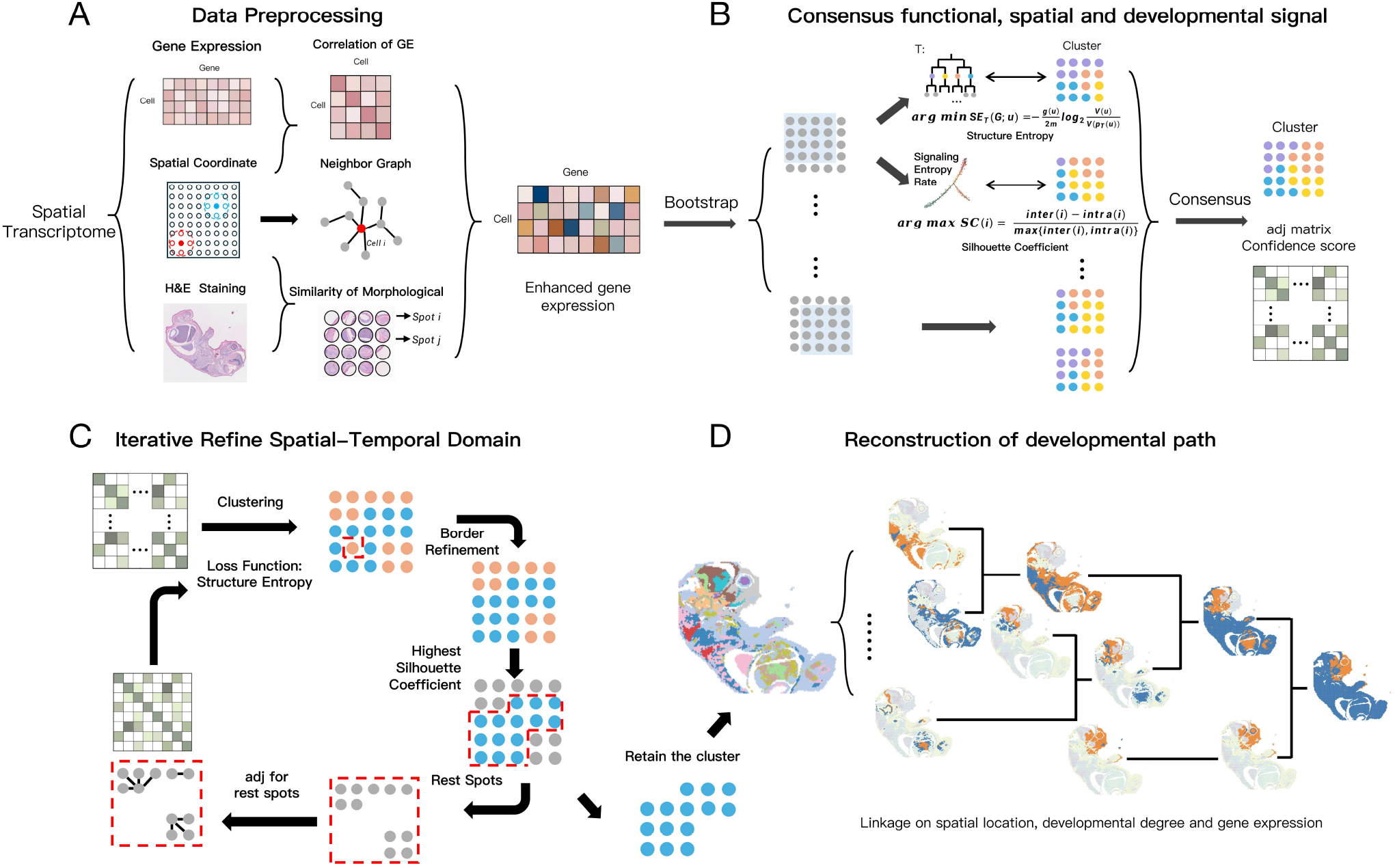
The flowchart of stTrace. (A). Data processing. We integrated gene expression, spatial location, and morphological information from Spatial transcriptomics data to generate enhanced gene expression; (B). Based on the enhanced gene expression, we did clustering to obtain a partition that best captured spatial, functional and development information respectively. The clustering that implies the overall structure from the aspect of gene expression similarity is obtained by minimizing the Structure Entropy. The development information is measured by the Signaling Entropy Rate, and we perform clustering to maximize the Silhouette Coefficient. The two types of signals are aggregated by bootstrap and consensus then generate the confidence score matrix; (C). Iterative refine spatial-temporal domain. For each iteration, the graph is constructed by taking the confidence score as the adjacency matrix. After applying clustering and border refinement, we retained the cluster with the highest Silhouette Coefficient score, which shows the best compactness at the developmental level. The remaining cells were processed for further clustering. The clustering in each iteration adopted Louvain taking the Structure Entropy as objective function, which aims to optimize the overall structure decomposition from gene expression similarity’s view. The iterations were repeated until all cells were clustered and we got spatial-temporal domains; (D). We did hierarchical clustering on the detected spatial-temporal domain and got the developmental path

Based on the varying resolutions of different spatial transcriptomics technologies, the enhanced expression matrix in our experiments was calculated at the spot level. For convenience in the text, we refered to each spot as a “cell”. Then, we used enhanced gene expression data as input to identify spatially continuous and functional similar regions by minimizing Structure Entropy. Additionally, we calculated the Signaling Entropy Rate for each cell as a measurement of developmental level and applied this to clustering by maximizing the Silhouette Coefficient to detect temporal continuous regions. By integrating these two clustering results, we obtained a confidence matrix (Fig. 2B). After that, given the confidence matrix as the similarity matrix, we conducted clustering using Louvain’s framework [4] while focusing on optimizing the Structure Entropy during each iteration (details on the clustering are described in Supplementary Materials). In addition, at each iterative step, we applied border refinement and calculated the Silhouette Coefficient aids to select the most promising domain and retain it. The final clustering results, referred to as spatial-temporal domains, have clearer regional boundaries, where cells within these regions exhibit similar biological functions and close developmental level (Fig. 2C). Subsequently, we performed hierarchical linkage on these domains to reveal the overall developmental paths of the samples (Fig. 2D). stTrace aims to identify hierarchically related spatial-temporal continuous regions by capturing the temporal, spatial and biological functional characteristics of cells, thereby revealing the developmental paths within the tissue.

### Detect functional similar and spatial continuous region by Structure Entropy

To recognize continuous spatial regions with similar gene expression, we performed clustering based on the enhanced gene expression by minimizing the Structure Entropy. Structural information theory quantifies the uncertainty inherent in the dynamics of a graph [11], while identifying partition in a graph that minimizes Structure Entropy essentially means a division that most accurately represents the original graph with the least random variations and noise[39, 6]. Here, the graph refers to the cell graph, more specifically, where edges refer to the association between cells based on their gene expression patterns, spatial coordinates, and corresponding H&E-stained images, which are calculated according to correlation on the enhanced gene expression. To calculate Structure Entropy, firstly, consider a coding tree *T* in the weighted cell graph *G*. Suppose the root *λ*_*T*_ represents the whole tissue, each node of *T* contains a set of cells. Bifurcate of the tree means the set of cells belonging to the typical tree node is partitioned by their child nodes, implying the region is divided into smaller regions. Then in the result *T*, each leaf node is a structural, functional, and spatial similar region, containing a set of cells. The coding tree *T* can be used to represent the hierarchical relationship between those regions. The cells represented by a node *u* ∈ *T* are shown as *b*_*T*_ (*u*), the parent node of node *u* is *p*_*T*_ (*u*), and the volume *V* (*u*) of this node is:

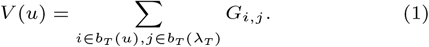

where *G*_*i,j*_ is the weighted graph generated by the correlation of enhanced gene expression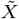 and filtering. The Structure Entropy of node *u* is defined as:

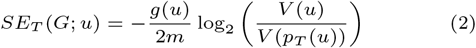

where

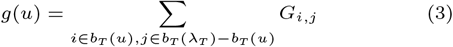

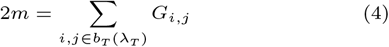

Denote the leaf node in *T* as *t*(*i*) where cell *i* belongs to, then the Structure Entropy of cell *i* is:

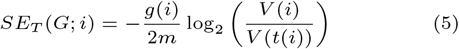

where

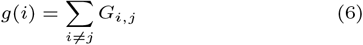

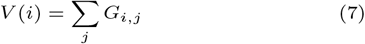

The Structure Entropy *SE*_*T*_ (*G*) of a coding tree *T* is the sum of the Structure Entropy of nodes *SE*_*T*_ (*G*; *u*) and cells *SE*_*T*_ (*G*; *i*):

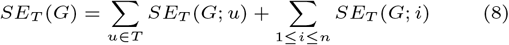

We identified the most cohesive partition of the graph following Eq.13, dividing cells into continuous spatial regions where cells display similar gene expression profiles, according to Supplementary Methods. This method ensures that the partition captures the intrinsic structure of cellular relationships and maintains the hierarchical organization between regions.

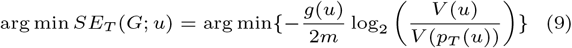

### Temporal continuous region from development degree

The temporal continuity of cells can be reflected by the closeness in their developmental level. We calculated cellular developmental level using the Signaling Entropy Rate, as proposed by Teschendorff and Enver [29, 30]. This method measures the diversity of signal transduction within protein-protein interaction networks based on the gene expression profiles in the transcriptome, thereby reflecting the differentiation potential of individual cells. We downloaded protein-protein interaction network from String database, denoted as *A*. Protein *m* and protein *n* are considered as connected when *A*_*mn*_ = 1, otherwise *A*_*mn*_ = 0. Then we convert *A* to weighted by considering gene expression data for each cell *i*, assuming that the weight of the edges *W*_*mn*_ between protein *m* and protein *n* is proportional to the gene expression level *X*_*m*_ and *X*_*n*_. Normalizing the sum of outgoing edge weights for each node to 1, defining a random walk on the network. This results in a matrix *P*,

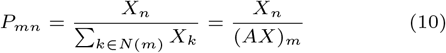

Where *N* (*m*) is the neighbor of protein *m*. The Signaling Entropy Rate can be defined as the entropy rate of cell *i* over the protein-protein interaction network,

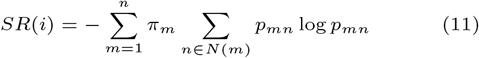

Where *π* is the steady-state distribution, which satisfies *πP* = *π* and the normalization condition *π*^*T*^ 1 = 1 [29].

The developmental level is regarded as the cell’s signal at the temporal aspect, where cells that are closer in developmental time will exhibit more similar gene expression patterns and cellular behaviors. We calculated the Signaling Entropy Rate for each cell and then the distance matrix, which quantifies the temporal distance between cells based on their developmental status. Using this matrix, we built a weighted graph where the edges represent the gene expression similarity between cells in the temporal aspect. Finally, we used the Silhouette Coefficient as the objective function to cluster cells according to their developmental level (Supplementary Methods).

Silhouette Coefficient [24] is an evaluation metric to assess clustering effectiveness, measuring the compactness and separation of cells within clusters. For each cell *i*, the Silhouette Coefficient *sc*(*i*) is given by the formula:

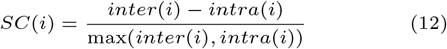

Where *intra*(*i*) is the average difference of developmental level *SR* between cell *i* and all other cells in the same cluster; *inter*(*i*)is the average difference of developmental level *SR* between cell *i* and all cells in the nearest cluster; When there is only one sample in the cluster, *SC*(*i*) is typically 0. The global Silhouette Coefficient *SC*(*global*) is the average of all individual Silhouette Coefficients *SC*(*i*), used to measure the overall quality of the clustering.

Cells were divided into several continuous regions along the temporal dimension with the objective function

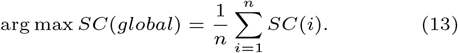

This clustering method ensures that cells within the same region not only exhibit similar gene expression profiles but also share comparable developmental stages. Through these continuous regions, dynamic changes in the cellular development process can be identified.

### Consensus signal of structure and development

To effectively integrate gene expression, spatial information, and developmental status, we combined the features obtained in the temporal, spatial and functional dimensions as mentioned above by bootstrap experiment. First, to ensure the robustness of the algorithm and the stability of the results, we repeated clustering 10 times. In each run, we randomly selected 80% of the cells and independently clustering the cells in both dimensions. Then we computed the probability of each pair of cells being assigned to the same cluster across the multiple runs and generated the confidence score matrix *C*. The confidence score represents the consistency of cell clustering outcomes across different trials and serves as a consensus signal that encapsulates the inherent relationships between cells in spatial, functional and temporal aspects. A higher confidence score indicates a strong likelihood that two cells have a higher possibility to be in the same cluster where cells are similar in gene expression, neighbor in space, and also close in development status. This matrix is then used as an adjacency matrix of the graph for further iterative refinement, ensuring that the final clusters reflect spatial continuity, biological functional and developmental similarity.

### Iterative refine spatial-temporal domain

Iterative refinement smooths the border and detects the most promising region that reflects the partition in temporal, spatial and functional aspects in each iteration, resulting in the spatial-temporal domains. First, a weighted graph *G*′ is constructed using confidence scores matrix *C*. Each cell connects to its top 30 neighboring cells ordered by the confidence scores. If cell *i* is one of the 30 nearest neighbors of cell *j*, or cell *j* is one of the 30 nearest neighbors of cell *i*, then *e*_*ij*_ equals the confidence score between the two cells; otherwise, *e*_*ij*_ is 0. Given graph *G*′, we clustered to get the partition that minimizes the Structure Entropy of *T*. The clustering method adopt the framework of Louvain [4], and its detail are described in the Supplementary. Then, we refined the cluster boundaries by examining the clustering distribution of neighboring cells. For a cell *i*, if 80% of cell *i*’s neighboring cells belong to cluster A, while cell *i* is assigned to cluster B, we reassign cell *i* to cluster A to align it with the majority of its neighbors. After border refinement, we computed the Silhouette Coefficient [24] on the cells’ developmental level within each region. The region with the highest Silhouette Coefficient score is the spatial continuous region with the highest compactness along the temporal aspect in this iteration. Therefore we retained the cluster with the highest score as a spatial-temporal domain, isolating them from the current graph *G*′, while continuing partition refinement for the resting cells. For the resting cells, we updated the *G*′ by renewing the neighboring cells according to confidence scores. Subsequently, clustering and border refinement is performed on the updated *G*′. The iteration ends when every cell has been assigned to a spatial-temporal domain.

### Reconstruction of developmental path

The spatial-temporal domains exhibit continuity in spatial, functional and temporal dimensions, reflecting the underlying developmental processes. This continuity region naturally contains hierarchical relationships between these domains as the development. Therefore, we performed hierarchical clustering on the identified temporal domains to reconstruct the developmental path. First, we get each domain’s first principal component as representative features. Based on these features, we constructed a correlation matrix reflecting the gene expression similarities between the domains. Second, regions that are spatially adjacent and share similar gene expression profiles tend to exhibit similar developmental progression, thus we calculated the minimum Euclidean distance between any two points from the two domains as spatial distance. Third, we computed the average Signaling Entropy Rate in each domain and generated the temporal distance matrix that reflects the developmental level differences between domains. We merged these three matrices, the gene expression distance, the spatial distance, and the developmental level distance, to form a linkage input matrix. Using this combined matrix, we applied the average linkage algorithm to generate a hierarchical clustering tree. This clustering tree would reveal the overall progression of cellular development.

### Application to real data and evaluation

We applied stTrace to mouse embryos and human breast cancer, which are public spatial transcriptome data from 10X. For each dataset, we identified spatial-temporal domains and developmental paths to show the application in both development and cancer metastasis. we compared domains detected by stTrace and other traditional spatial domain detection methods, including Seurat, stLearn, DeepST, stDyer [38], GraphST [12], CellCharter [34] and STAGATE. More specifically, to evaluate the performance of stTrace, we did the analysis in the following four aspects.

To assess the continuity of spatial-temporal domains identified by stTrace, we evaluated both spatial and temporal aspects. In the spatial dimension, we examined whether the detected domains correspond well to HE-stained tissue morphology and display clear patterns in UMAP-reduced space. Since stTrace is designed to detect spatial-temporal domains and reconstruct developmental trajectories, which is different from identify spatial domains, traditional metrics such as Adjusted Rand Index (ARI) are not suitable, especially given the absence of ground truth spatial annotations in developmental datasets like embryos and tumors. In the temporal dimension, no gold standard currently exists for spatial-temporal domains. To address this gap, we employed indirect indicators of developmental progression, including Signaling Entropy Rate, Pseudotime, and RNA velocity. We further introduced two novel metrics—inter-difference and intra-difference—to quantitatively measure domain separation and consistency in developmental dimension. Inter-difference reflects the disparity in average developmental level between domains, while intra-difference quantifies the deviation of individual cells from the average level of its assigned domain. An ideal clustering yields a larger inter-difference (indicating clear developmental separation between domains) and a smaller intra-difference (indicating developmental consistency within each domain). Statistical significance of performance improvements was assessed via pairwise Mann-Whitney U tests between stTrace and comparative methods. Additionally, we evaluated spatial compactness and separation by computing intra- and inter-domain differences in terms of spatial distances based on the coordinates in UMAP-reduced latent space.

Second, we validate the detected developmental path using different measurements that reflect the development level and check the consistency. The measurements include Signaling Entropy Rate [29, 30], identifying the developmental status of cells based on gene expression profiles, Pseudotime[22], and Single Cell Velocity [2] focusing on dynamic changes in the temporal dimension.

Third, by analyzing the functional annotation referring to the domain and path detected, we showed that stTrace can provide biologically meaningful information. we compared its results with MOSTA [5] in mouse embryos and BayesSpace [40] in human breast cancer. The result shows our method can infer the directional spread of invasive cancer cells mapped with both cellular developmental levels and gene expression trends.

Fourth, to evaluate the necessity of incorporating cellular developmental information in spatial-temporal domain identification, we performed an ablation study by comparing the clustering performance with and without integrating developmental progression. In the consensus signal of structure and development module, we focused on the regions with similar function and spatially continuous which detected by Structure Entropy, without considering the continuous in cell developmental. During the iterative refinement of spatial-temporal domains, we optimized the identified regions by evaluating the Silhouette Coefficient based on Euclidean distances between cells, rather than developmental levels.

## Results

### Spatial-temporal domain has higher resolution and better compactness in developmental level

To evaluate stTrace, we compare stTrace with Seuart, stLearn, DeepST, STAGATE, stDyer, GraphST and CellCharter in the mouse embryo and human breast cancer. The result (Fig. 3 and Fig. 4) shows the identified spatial-temporal domains exhibited higher resolution, and demonstrates continuity in spatial, functional and temporal dimensions. Fig. 3A and Fig. 4A demonstrate the comparison on the detected domains of different methods and H&E-stained images on mouse and human data. The images in red and blue boxes highlight differences in domains identified by various methods within specific complex tissue regions. The result of DeepST, GraphST and CellCharter look coarse, and these domains cover larger areas. Although these methods can capture general regional divisions, they are limited in resolution, which may be important for detailed structural analysis. Although Seurat, stLearn, stDyer and STAGATE offer more refined distinctions compared to DeepST, GraphST and CellCharter, identifying more domains, their ability to resolve fine-grained structures is limited to only certain regions in both the mouse and human datasets. stTrace outperforms in complex tissue structures that require higher resolution, such as the mouse retina region in the mouse embryo, and further division inside invasive cancer regions. The spatial-temporal domains detected by stTrace are well-mapped to the distribution observed in H&E-stained images.

**Fig. 3.**
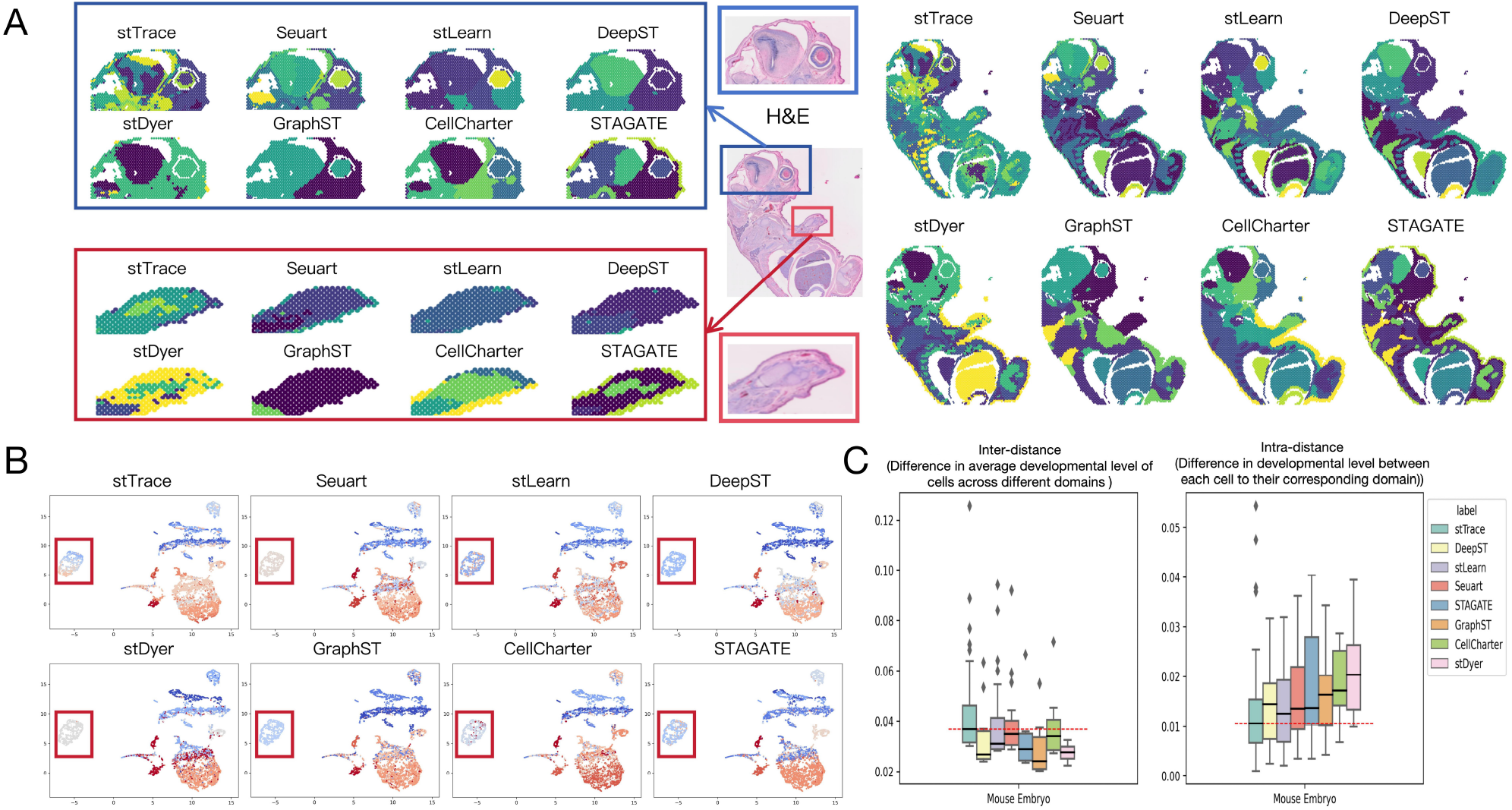
Comparison of stTrace with other spatial domain identification methods on Mouse Embryo data. (A) Spatial-temporal domains identified by stTrace (integrating developmental and expression data), stTrace (SE, based on expression only), and other methods including Seurat, stLearn, DeepST, stDyer, GraphST, CellCharter, and STAGATE. Enlarged views highlight structural differences (red/blue boxes). (B) UMAP plots showing domain distributions colored by average developmental level (blue: early stage; red: late stage). (C) The left sub-figure shows the inter-difference, i.e., the difference in average developmental levels across domains; the right sub-figure shows the intra-difference, i.e., the deviation of each cell’s developmental level from its domain’s average. Higher inter-difference and lower intra-difference indicate better domain separation and internal consistency, red line denotes stTrace median.

**Fig. 4.**
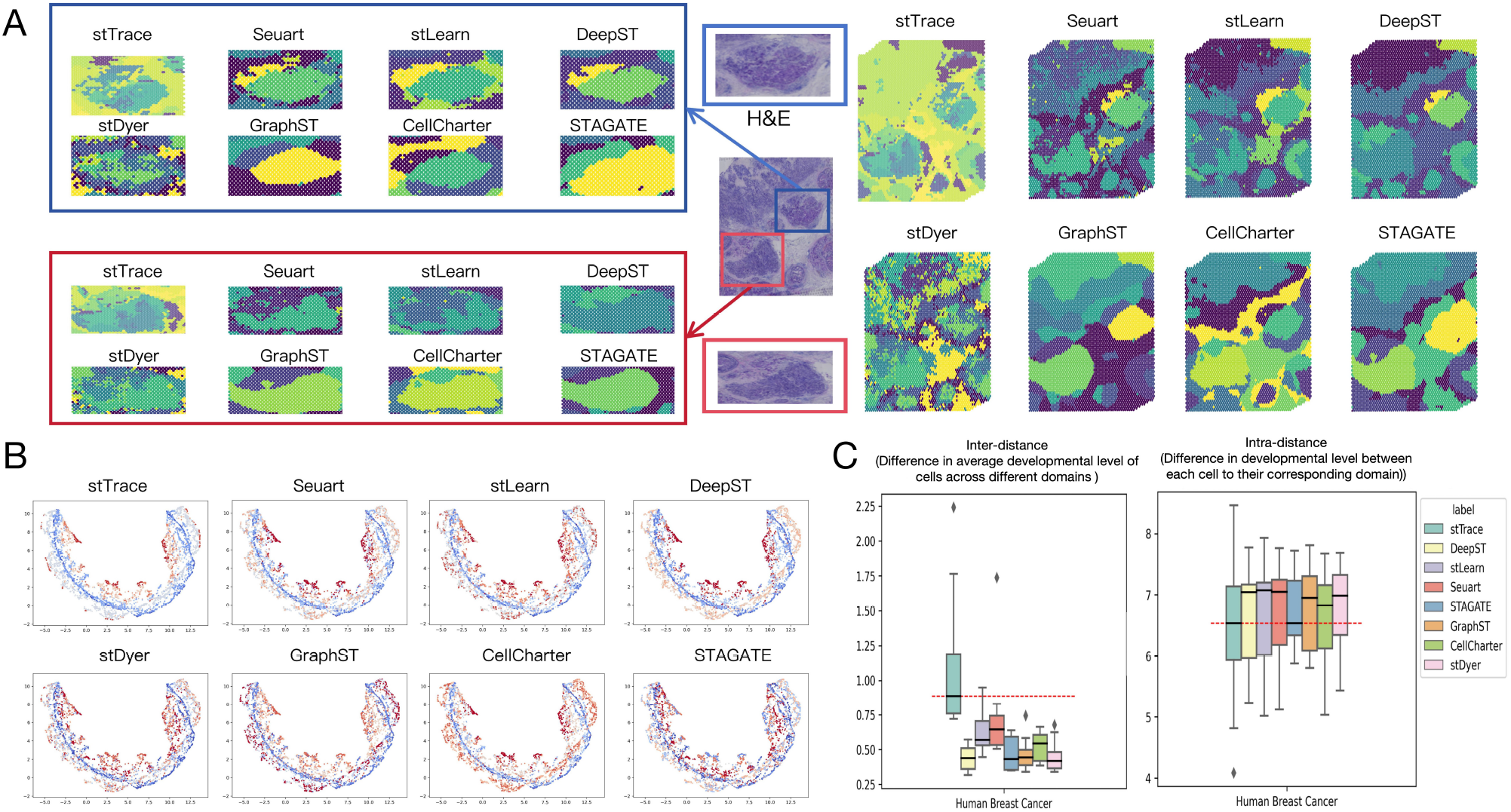
Comparison of stTrace with other spatial domain identification methods on Human Breast Cancer data. (A) Spatial-temporal domains identified by stTrace (integrating developmental and expression data), stTrace (SE, based on expression only), and other methods including Seurat, stLearn, DeepST, stDyer, GraphST, CellCharter, and STAGATE. Enlarged views highlight structural differences (red/blue boxes). (B) UMAP plots showing domain distributions colored by average developmental level (blue: early stage; red: late stage). (C) The left sub-figure shows the inter-difference, i.e., the difference in average developmental levels across domains; the right sub-figure shows the intra-difference, i.e., the deviation of each cell’s developmental level from its domain’s average. Higher inter-difference and lower intra-difference indicate better domain separation and internal consistency, red line denotes stTrace median.

Additionally, we plot domains obtained from different methods in UMAP space [15], and compared the distribution of cell developmental level, as shown in Fig. 3B and Fig. 4B. The color represents the average developmental level of cells for each domain. First, Seuart, stLearn, DeepST, STAGATE, stDyer, GraphST and CellCharter have areas mixing points with different clusters, with apparent color variation between adjacent cells. In contrast, in stTrace, the partition is more distinct in the UMAP space, which implies that stTrace has partitions that better fit the feature captured in the UMAP space. Second, the color distribution in stTrace shows a smoother and clearer color gradient, from blue to red. This gradient reflects the cellular development continuity inside the domain and the changes across domains are well captured in stTrace. As a comparison, the color transition of other methods is more vague and has less correlation with developmental stages. Third, the stTrace generates partitions with higher resolution. For example, in the area indicated by the red squares in Fig. 3B, stTrace is the only method that divides the points according to the development level. Seurat, DeepST, stDyer and GraphST treat them as one cluster without further distinguishing, while partition in stLearn does not capture the difference in developmental level.

Spatial-temporal domains should exhibit continuity in spatial, functional and temporal dimensions, meaning that cells within the same region are not only spatially adjacent with similar gene expression but also at similar developmental stages. The idea of spatial-temporal domain is novel and currently the gold-standard for this kind domain is lacking. To evaluate the compactness for the domain detected by stTrace, we calculated the domains’ inter-difference and intra-difference and compared to Seurat, stLearn, DeepST, stDyer, GraphST, CellCharter and STAGATE, as described in the method section. The left sub-figure of Fig. 3C and Fig. 4C is the boxplot of cell’s inter-distance across all detected domains for different methods, while the right sub-figure of Fig. 3C and Fig. 4C is the boxplot of intra-difference for each domain, representing the difference between the developmental level of each cell and the mean developmental level of the corresponding domain. A greater distance between domains signifies a more compact partitioning, and smaller intra-difference indicates greater consistency in the developmental level of the cells within the domain. In both datasets, the inter-difference identified by stTrace is larger than that identified by other methods, and the intra-difference is smaller than other methods. We use the Mann-Whitney U test to assess whether the increase in inter-difference and decrease in intra-difference are statistically significant. The results are shown in Table.1 for Mouse Embryo data and Table.2 for Human Breast Cancer data. In the Mouse Embryo dataset, stTrace achieved significantly lower intra-difference compared to several methods, including DeepST (p = 0.018), stLearn (p = 0.031), STAGATE (p = 0.012), GraphST (p = 0.005), and stDyer (p = 0.002), indicating stronger developmental consistency within domains. For inter-difference, although none of the comparisons reached strong statistical significance (p *<* 0.05), CellCharter showed a marginally significant result (p = 0.020), suggesting a slight advantage in domain separation. In the Human Breast Cancer dataset, inter-difference values achieved by stTrace were significantly higher than those by all other methods (p *<* 0.05), demonstrating better separation between domains. For intra-difference, stTrace significantly outperformed STAGATE (p = 5.94e-07), GraphST (p = 5.83e-03), and stDyer (p = 6.39e-04), confirming the method’s effectiveness in maintaining developmental consistency within domains.

**Table 1.** Mann-Whitney U Test Results in Mouse Embryo Data.(stTrace VS others)

| p-value | DeepST | stLearn | Seurat | STAGATE | GraphST | CellCharter | stDyer |
| --- | --- | --- | --- | --- | --- | --- | --- |
| **inter-difference** | 3.9985e-01 | 4.5267e-01 | 9.4330e-02 | 1.2812e-01 | 1.2976e-01 | **2.0051e-02** | 5.5109e-02 |
| **intra-difference** | **1.8140e-02** | **3.0903e-02** | 3.2993e-01 | **1.1841e-02** | **4.5547e-03** | 1.5950e-01 | **1.6588e-03** |

**Table 2.** Mann-Whitney U Test Results in Human Breast Cancer Data.(stTrace VS others)

| p-value | DeepST | stLearn | Seuart | STAGATE | GraphST | CellCharter | stDyer |
| --- | --- | --- | --- | --- | --- | --- | --- |
| inter-difference | 5.8358e-03 | 1.1925e-02 | 9.8995e-03 | 6.9938e-04 | 2.5913e-03 | 2.5913e-03 | 2.0321e-03 |
| intra-difference | 2.9799e-01 | 7.3744e-01 | 6.8505e-01 | 5.9420e-07 | 5.8323e-03 | 2.2120e-01 | 6.3928e-04 |

Besides, Supplementary Fig. S1 compares the intra- and inter-differences in Euclidean distances based on UMAP-space, further validating the spatial and functional compactness and separation achieved by stTrace. The figures show that the spatial-temporal domains identified by stTrace exhibit better consistency in terms of developmental level, indicating that these domains maintain continuity along the temporal dimension. These results collectively validate that stTrace achieves better performance in terms of separation between domains and consistency within domains.

### stTrace identifies developmental and invasion direction across spatial-temporal domains

stTrace reconstructs the hierarchical relationship among spatial-temporal domains, which can reveal the developmental paths within the Mouse Embryo tissue. In the mouse embryo data, from cell type annotation of spatial-temporal domains detected by stTrace (Fig. 2.5A), as well as the Single Cell Velocity [2] direction across domains (Fig. 2.5B), we observed that brain show the highest maturity, followed by abdominal organs, limbs, and the trunk. The inferred paths are aligning with the developmental timeline established by MOSTA [5]. To further validate the inferred trajectory, we plotted distributions of cells’ developmental level in space, which are Signaling Entropy Rate [29, 30], Pseudotime[22], and Single Cell Velocity of cells in Fig. 5C. The distribution of cells’ developmental level can be well mapped with the spatial-temporal hierarchical structure identified by stTrace. The developmental tree constructed based on the hierarchical relationships of spatial-temporal domains shown in Supplementary Fig. S3.

**Fig. 5.**
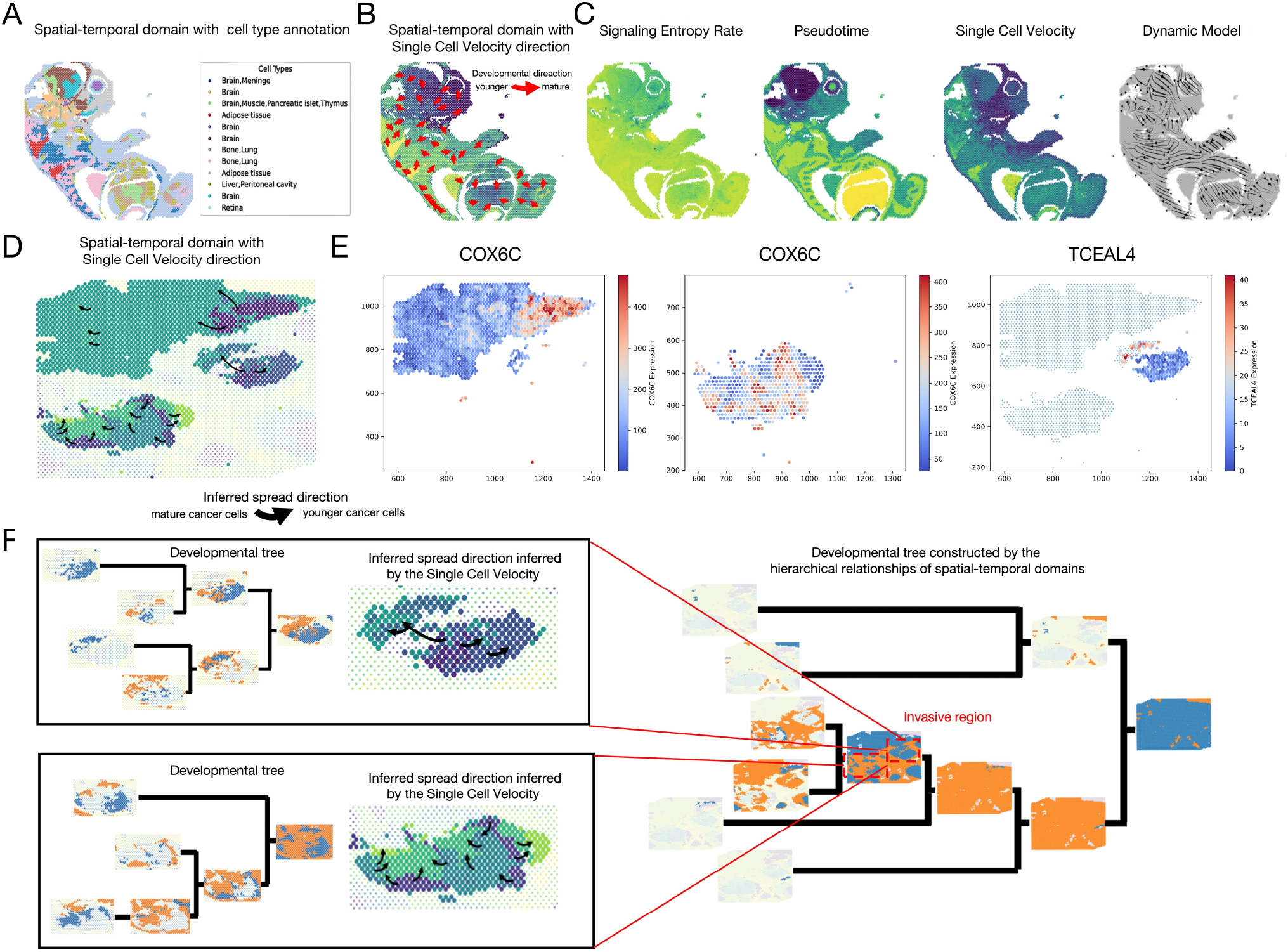
Developmental path constructed with hierarchical relationships among spatial-temporal domains detected by stTrace. **(A–C) Results from mouse embryo data**. (A). Cell type annotation within the identified spatial-temporal domains;(B). Developmental direction inferred by the Signaling Entropy Rate. The figure uses color coding to represent the average developmental level of each domain, with the gradient from light to dark indicating the progression from younger to more mature stages. Arrows indicate the direction of cell development, showing the trajectory from younger to more mature cells. The brain cells are in the most mature developmental stage, followed by the abdominal organs, limbs, and trunk, aligning with the organ developmental timeline obtained from MOSTA analysis;(C). Distribution of Signaling Entropy Rate, Pseudotime, Single Cell Velocity (latent time) across cells, and the dynamic model generated based on latent time, darker colors represent higher developmental level, reflecting the change in developmental level. The maturity distribution of spatial-temporal domains identified by stTrace closely matches the developmental trends observed; **(D–F) Results from human breast cancer data**. (D) Spread path of invasive cancer cells in Human Breast Cancer. Average Signaling Entropy Rate calculated for each spatial-temporal domain, with darker colors indicating more mature developmental levels. Black arrows represent the inferred direction of cancer cell spread, pointing from more mature regions (indicative of advanced stages of cancer) towards newer, less developed areas in the invasion zone; (E) Expression distribution of cancer-related genes COX6C and TCEAL4 identified from differential expression analysis across the three invasion regions. The expression levels of these genes show distinct patterns along the inferred cancer cell spread direction; (F). Developmental tree constructed based on the hierarchical relationships of spatial-temporal domains. This tree visualizes the developmental path of cancer cells across the human breast cancer sample, along with the inferred spread direction of cancer cells within the invasive regions.

We also apply stTrace to breast cancer data, the result shows it can identify high-resolution domains within each invasion zone, and capturing the temporal evolution of cancer progression. Based on the detected spatial-temporal domain, we calculated the average Signaling Entropy Rate to denote the cancer development level for each domain. The inferred spread direction is shown in Fig. 5D, where black arrows indicate the inferred spread direction, which points from more developed areas to less developed areas. Then we conducted differential expression across domains within each invasion zone and listed the identified genes in Supplementary Tab.S2-S4. Because domains in each invasion zone correspond to different stages of invasion development, the genes detected by differential expression across spatial-temporal domains may be related to cancer invasion.

We plot the expression distribution for the detected genes in Fig. 5E and Supplementary Fig. S4, and marker genes for cancer invasion in Supplementary Fig. S2. The marker genes, obtained from a previous publication [40], includes the proliferation marker gene MKI67 [33], genes encoding the oncogene ERBB2 [9], pro-apoptotic protein BAX [10], and tumor suppressor gene PTEN [27]. These genes’ expression abundance changes shown in the distribution can be well mapped to the inferred spread direction by stTrace. For instance, ERBB2 expression increases along the invasion path, consistent with its role in promoting tumor invasiveness[9]. Fig. 5F illustrates the developmental path across the human breast cancer sample, as well as the spread directions of cancer cells within the invasive region. These results demonstrate that stTrace can effectively integrate developmental and transcriptomic information to infer cancer spread directions and key genes.

### Incorporating developmental information enhances spatial-temporal domain identification

Incorporating cellular developmental information leads to more coherent spatial-temporal domains, both in spatial and time dimension. Ablation studies show that, compared to stTrace(SE) which detect domains without temporal information, stTrace identifies more distinct and spatially continuous domains, separating biologically adjacent regions such as retina and bone in mouse embryo and tumor-infiltrated areas in human breast cancer (Fig. 6A,D). UMAP visualization reveals stTrace has smoother developmental gradients and greater intra-domain consistency (Fig. 6B,E). Quantitative metrics show stTrace higher inter-difference in mouse embryo and lower intra-difference in breast cancer data (Fig. 6C,F), confirming that considering developmental information in stTrace can improve compactness of the detected domain.

**Fig. 6.**
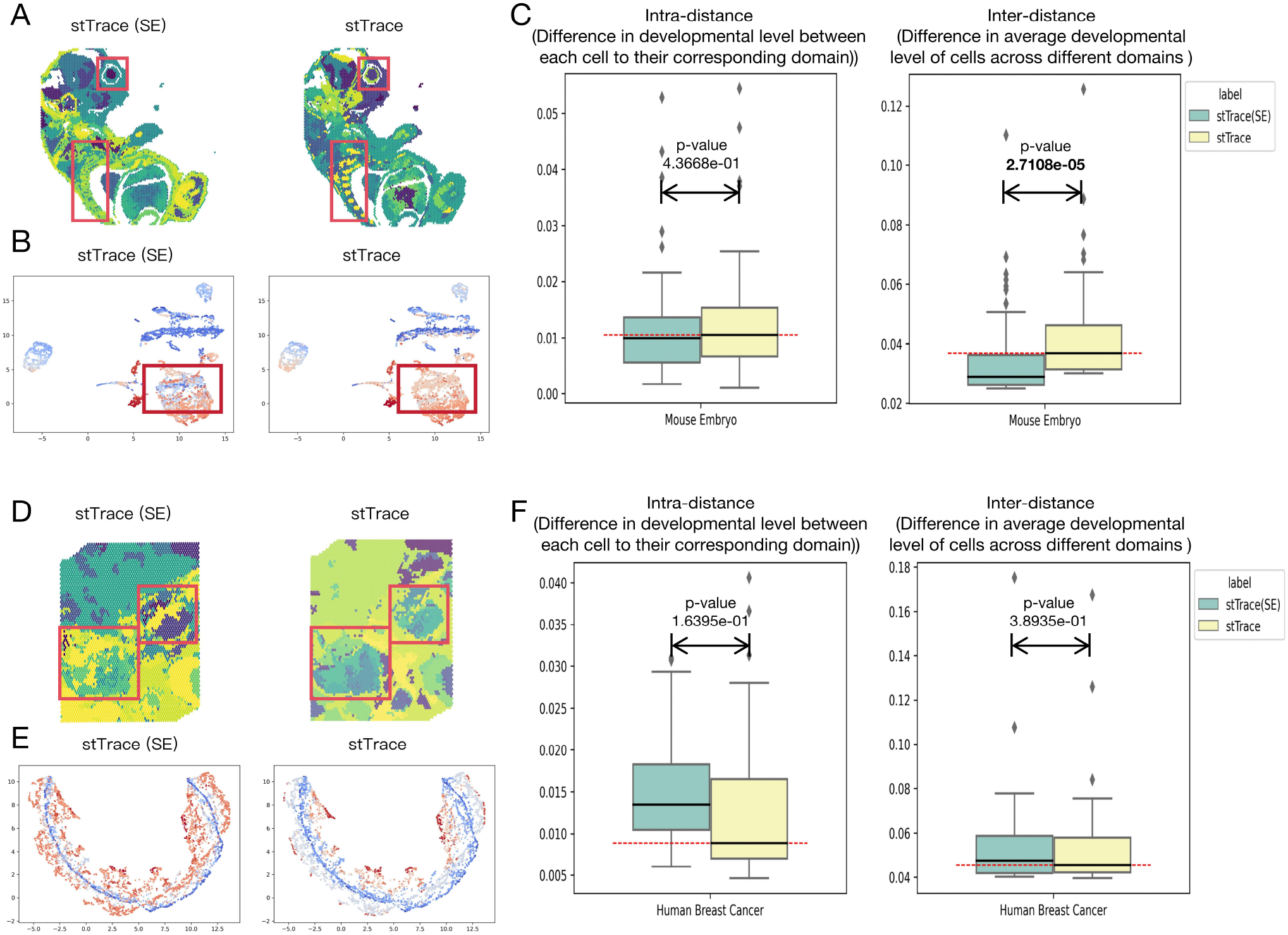
Ablation Study on Spatial-Temporal Domain Identification, regions marked by red boxes indicate areas with differences between the two methods. (A, D) The results demonstrate the impact of excluding or incorporating cellular developmental information on spatial domain coherence. Excluding developmental progression results in fragmented spatial domains, while incorporating developmental information enhances the spatial coherence of the identified domains; (B, E) UMAP embeddings of the clustering results, colored by developmental levels, show a gradual color transition in domains identified with developmental information, indicating consistent progression along the developmental timeline. Cells within each domain exhibit similar developmental levels, reflecting stronger biological consistency and internal homogeneity; (C, F) Quantitative evaluation of clustering performance through inter-difference and intra-difference metrics, we are expecting larger inter-difference and smaller intra-difference. In the mouse embryo dataset (C), incorporating developmental information significantly increases the inter-difference, enhancing the distinction between spatial-temporal domains. In contrast, in the human breast cancer dataset (F), the inclusion of developmental progression reduces the intra-difference, indicating improved consistency within each domain.

stTrace demonstrates robust adaptability to diverse developmental metrics, consistently reconstructing high-resolution spatial-temporal domains across different representations of cellular progression. By replacing the Signaling Entropy Rate [29, 30] with alternative metrics such as Pseudotime [22] and Single Cell Velocity (latent time) [2], we reconstructed the developmental trajectories of mouse embryos within the stTrace framework. Each metric emphasizes a distinct aspect of developmental biology: Signaling Entropy Rate reflects a cell’s differentiation potential based on transcriptomic heterogeneity; Pseudotime estimates a cell’s relative position along a developmental path; and Single Cell Velocity captures dynamic gene expression changes to infer temporal progression. Despite these conceptual differences, all three metrics provide temporal information during development. The result of the three metrics is consistent in developmental hierarchies. As shown in Supplementary Fig. S6, three metrics all indicate that brain cells emerge as the most mature, followed by those in abdominal organs, limbs, and trunk. The resulting lineage trees consistently show the same merging order, from trunk to abdomen and then to brain, indicating a stable and biologically meaningful developmental path. In addition, stTrace that adopt three different developmental information all reveal fine-grained spatial-temporal domains with higher resolution than traditional methods. These results suggest that stTrace is adaptable to diverse measures of cellular progression or development trajectory, and is robust to the choice of developmental metric.

## Conclusion

In this work, we propose stTrace, a novel algorithm that integrates gene expression, spatial information, and developmental level to identify spatial-temporal domains and reconstruct developmental paths. By combining structure entropy, signaling entropy rate, and silhouette coefficient, stTrace ensures spatial continuity, functional similarity, and temporal consistency. It outperforms existing methods in capturing fine-grained spatial structures and developmental transitions. Applied to mouse embryo and human breast cancer data, the identified domains show higher resolution and developmental coherence, and the reconstructed trajectories align well with Pseudotime [22], Single Cell Velocity [2], and MOSTA[5].

## Author contributions statement

Jiaxing Chen conceived the study, designed the algorithm, guided the experiment, and refined the manuscript. Zhangdi Song designed and implemented stTrace, conducted experiments and wrote the article. Changyu Zheng did experiments and visualization. All authors read and approved the final manuscript.

## Acknowledgments

This work is supported by the National Natural Science Foundation of China (Grant No. 32200526) with match fund provided by UIC Research Grant UICR0300014, and the Guangdong Provincial/Zhuhai Key Laboratory IRADS (2022B1212010006) and Guangdong Higher Education Upgrading Plan (2021-2025) with UIC Research Grant UICR0400025-21, and UIC Start Up Fund UICR0700039-22.

