## Supplemental methods and figures for "stTrace: Detecting Spatial-Temporal Domains from spatial transcriptome to Trace Developmental Path"

*Journal Title Here*, 2022, pp. 1–8

doi: DOI HERE
Advance Access Publication Date: Day Month Year
Paper

PAPER

### stTrace: Detecting Spatial-Temporal Domains from spatial transcriptome to Trace Developmental Path

Zhangdi Song,[1,2] Changyu Zheng[1] and Jiaxing Chen[1,*]

[1]Guangdong Provincial/Zhuhai Key Laboratory IRADS, Beijing Normal-Hong Kong Baptist University, 2000 Jintong Road, Tangjiawan, Zhuhai, 519087, Guangdong Province, China and [2]Hong Kong Baptist University, Kowloon Tong Kowloon, 100084, Hong Kong, China
FOR PUBLISHER ONLY Received on Date Month Year; revised on Date Month Year; accepted on Date Month Year

#### Abstract

#### Supplementary Materials

##### Data preprocessing

During the data preprocessing stage, we integrated the gene expression matrix, H&E-stained images, and spatial coordinates into an enhanced expression matrix, ensuring that spatial and gene expression information is comprehensively combined throughout the analysis, which is adapted from the framework in DeepST [4]. The enhanced gene expression data consider the curated correlation between cells with their neighboring cells in terms of both gene expression and morphology, where the neighboring cells are selected based on spatial locations.

For spatial transcriptome data with morphological information, first, we segment the images (H&E-stained slices) with the coordinates of each cell to obtain the partial images, which will be transformed and enhanced by using torchvision.transforms() function. With the convolutional neural network model, we can extract the features of shape for each cell. Then we applied the principal component to the obtained features and selected the first 50 principal components as latent characteristics. After that, we calculated the correlation of spatial gene expression $X_{ij}$ between every two adjacent cells $i$ and $j$, the similarity of morphological $MS_{ij}$ between cell $i$ and adjacent cell $j$ can be calculated with cosine distance [4].

Meanwhile, we selected the neighboring cells based on the spatial coordinates. For a given cell $i$, we first calculated the Euclidean distance $d_{ik}$ between $i$ and all other cells. Then, we selected top 5 closest neighboring cells for cell $i$, the distances are: $d_{i1}, d_{i2}, ...d_{i5}$. After identifying the five closest distances for all the cells, we calculated mean $\mu$ and variance $\sigma^2$:

$$\mu = \frac{1}{n}\sum_{i=1}^{n}\sum_{k=1}^{5} d_{ik} \tag{1}$$

$$\sigma^2 = \frac{1}{n}\sum_{i=1}^{n}\sum_{k=1}^{5}(d_{ik}-\mu)^2 \tag{2}$$

Next, the neighbor distance threshold $\gamma$ is defined as the sum of the mean and variance of the nearest distances:

$$\gamma = \mu + \sigma^2 \tag{3}$$

The cell $j$ will be considered as the neighbor cell of $i$ if and only if the distance between two cells $d_{ij}$ is less than $\gamma$. If cell $j$ is the neighbor of cell $i$, then spatial weight $SW_{ij} = 0$, otherwise $SW_{ij} = 1$.

With the correlation of gene expression, and similarity of morphological and spatial weight, the enhanced gene expression $\tilde{X}(i)$ could be denoted as:

$$\tilde{X}(i) = X_i + \frac{\sum_{j=1}^{n} X_j \cdot MS_{ij} \cdot X_{ij} \cdot SW_{ij}}{n} \tag{4}$$

where $X_i$ and $X_j$ are the gene expression for cell $i$ and adjacent cell $j$.

##### Materials and Methods

In this study, we adopt the Louvain algorithm [1] to achieve cell clustering based on spatial, functional and temporal dimensions. The traditional Louvain algorithm relies on modularity maximization to identify community structures within a network and requires a predefined number of iterations. In contrast, our method introduces the new objective function and iteration termination criteria.

We construct a graph based on the correlation matrix of cells, where each cell is represented as a node. For each cell, we identify its 30 most similar neighboring cells. Initially, each node is treated as an independent community, the number of initial communities equals the number of nodes. The algorithm then iteratively traverses each node, attempting to move it into the community of its neighboring nodes, while recording changes in the objective function after each operation. At the end of each traversal, the two nodes with the best optimization effect are merged into a new community, which is treated as a "super-node" that participates in subsequent iterations. This process continues until the objective function stabilizes.

© The Author 2022. Published by Oxford University Press. All rights reserved. For permissions, please

**Table 1.** The Calinski-Harabasz score for clustering results with different weights.

| Weight spatial | 0.1 | 0.2 | 0.3 | 0.4 | 0.5 | 0.6 | 0.7 | 0.8 | 0.9 |
|---|---|---|---|---|---|---|---|---|---|
| Calinski Harabasz Score | 285.5 | 285.5 | 275.2 | 262.7 | 258.7 | 260.5 | 175.2 | 295.1 | 204.2 |

##### *Detect functional similar and spatial continuous region by Structure Entropy*

We constructed a correlation matrix between cells with enhanced gene expression data, allowing us to extract spatial dimensional features of each cell from this matrix. During the clustering process, we selected Structure Entropy [5, 2] as the objective function to assess the quality of community division. A lower Structure Entropy value indicates a better clustering outcome.

Each cluster can be seen as a coding tree $T$, each node of $T$ contains a set of cells, the root $\lambda_T$ represents the whole cluster. The bins of each tree node are partitioned by their child nodes. The cells represented by a node $u \in T$ can be shown as $b_T(u)$, the parent node of node $u$ is $p_T(u)$, and the volume $V(u)$ of this node is:

$$V(u) = \sum_{i \in b_T(u), j \in b_T(\lambda_T)} G_{i,j}. \tag{5}$$

where $G_{i,j}$ is the weighted graph generated by the correlation of enhanced gene expression $\tilde{X}$ and filtering. The Structure Entropy of node $u$ is defined as:

$$S_T(G; u) = -\frac{g(u)}{2m} \log_2 \left( \frac{V(u)}{V(p_T(u))} \right) \tag{6}$$

where

$$g(u) = \sum_{i \in b_T(u), j \in b_T(\lambda_T) - b_T(u)} G_{i,j} \tag{7}$$

$$2m = \sum_{i,j \in b_T(\lambda_T)} G_{i,j} \tag{8}$$

Denote the leaf node in $T$ as $t(i)$ where cell $i$ belongs to, then the Structure Entropy of cell $i$ is:

$$S_T(G; i) = -\frac{g(i)}{2m} \log_2 \left( \frac{V(i)}{V(t(i))} \right) \tag{9}$$

where

$$g(i) = \sum_{i \neq j} G_{i,j} \tag{10}$$

$$V(i) = \sum_{j} G_{i,j} \tag{11}$$

The Structure Entropy $S_T(G)$ of a coding tree $T$ is the sum of the Structure Entropy of nodes $S_T(G; u)$ and cells $S_T(G; i)$:

$$S_T(G) = \sum_{u \in T} S_T(G; u) + \sum_{1 \leq i \leq n} S_T(G; i) \tag{12}$$

To ensure the robustness of the algorithm and the stability of the results, we randomly selected 80% of the cell nodes in each clustering experiment, repeating the experiment 10 times. After these 10 trials, we calculated the frequency of cell pair merges across experiments, producing a ”spatial confidence matrix” that reflects the likelihood of each pair of cells being grouped together in the spatial dimension.

##### *Temporal continuous region from development degree*

Similar to the last section, in the temporal dimension, we treat the developmental level of each single cell as a ”distance” in the time dimension, where each cell's developmental state is considered a relative time point. The differences in these developmental levels are transformed into a distance matrix, which serves as input for the algorithm. During the clustering process, we use the Silhouette Coefficient [3] as the objective function to assess clustering quality. A higher Silhouette Coefficient score indicates better clustering performance, meaning that the similarity within a community is higher and the similarity to other communities is lower.

For each cell $i$, the Silhouette Coefficient $sc(i)$ is given by the formula:

$$sc(i) = \frac{inter(i) - intra(i)}{\max(inter(i), intra(i))} \tag{13}$$

Where $intra(i)$ is the average distance between celll $i$ and all other cells in the same cluster; $inter(i)$is the average distance between celll $i$ and all cells in the nearest cluster; When there is only one sample in the cluster, $sc(i)$ is typically 0. The global Silhouette Coefficient is the average of all individual Silhouette Coefficients $sc(i)$, used to measure the overall quality of the clustering.

After 10 independent experiments, we can track the merging behavior of cells across each trial in the temporal dimension, constructing a ”temporal confidence matrix” that reflects the clustering structure of cells along the time series.

##### *Iterative refine Spatial-Temporal domain*

To effectively integrate spatial and temporal features, we designed a weighted coefficient to combine two confidence matrices, allowing both factors to be considered in the spatiotemporal clustering process. Experimental results demonstrated that the clustering performance is optimal when the spatial dimension is weighted at 20% and the temporal dimension at 80% (as shown in Tab.1). The combined confidence matrix is used as input for the algorithm, and Structure Entropy (Equation12) is employed as the objective function for cell clustering.

After completing the algorithm's iterations, we evaluate the clustering quality using the Silhouette Coefficient (Equation13). By comparing the Silhouette Coefficient values of each cluster, we select the group with the best clustering performance and retain it as the final spatiotemporal domain result. For the remaining unmerged cells, we perform re-clustering to ensure all cells are correctly assigned to their respective spatiotemporal domains, ultimately achieving a comprehensive spatiotemporal clustering result for all cells.

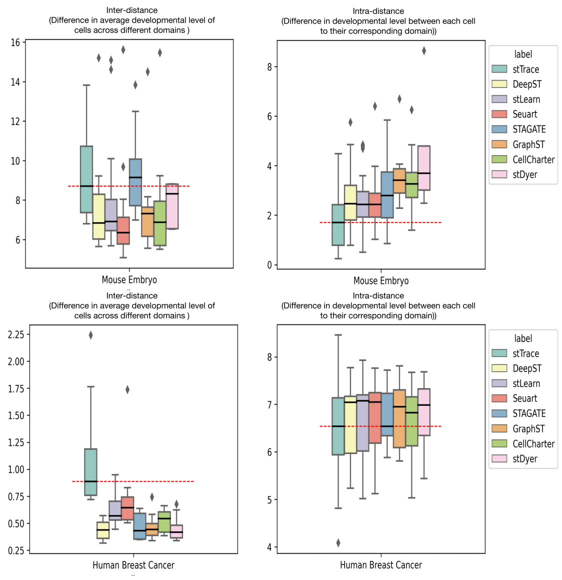


**Fig. S1. The Left sub-figures show the difference in Euclidean distances based on UMAP-reduced coordinates across different domains, which refers to inter-difference; The right sub-figures show the difference in Euclidean distances between each cell to the corresponding domain's centroid, which refers to intra-difference.**

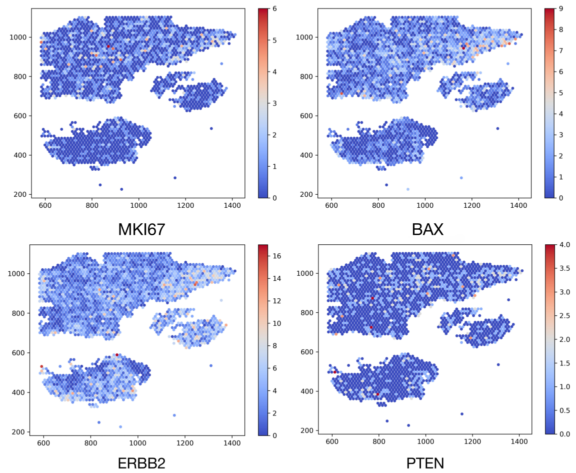


**Fig. S2. Expression distribution of key cancer invasion markers in the human breast cancer dataset, including the proliferation marker MKI67, the oncogene ERBB2, the pro-apoptotic protein BAX, and the tumor suppressor gene PTEN.**

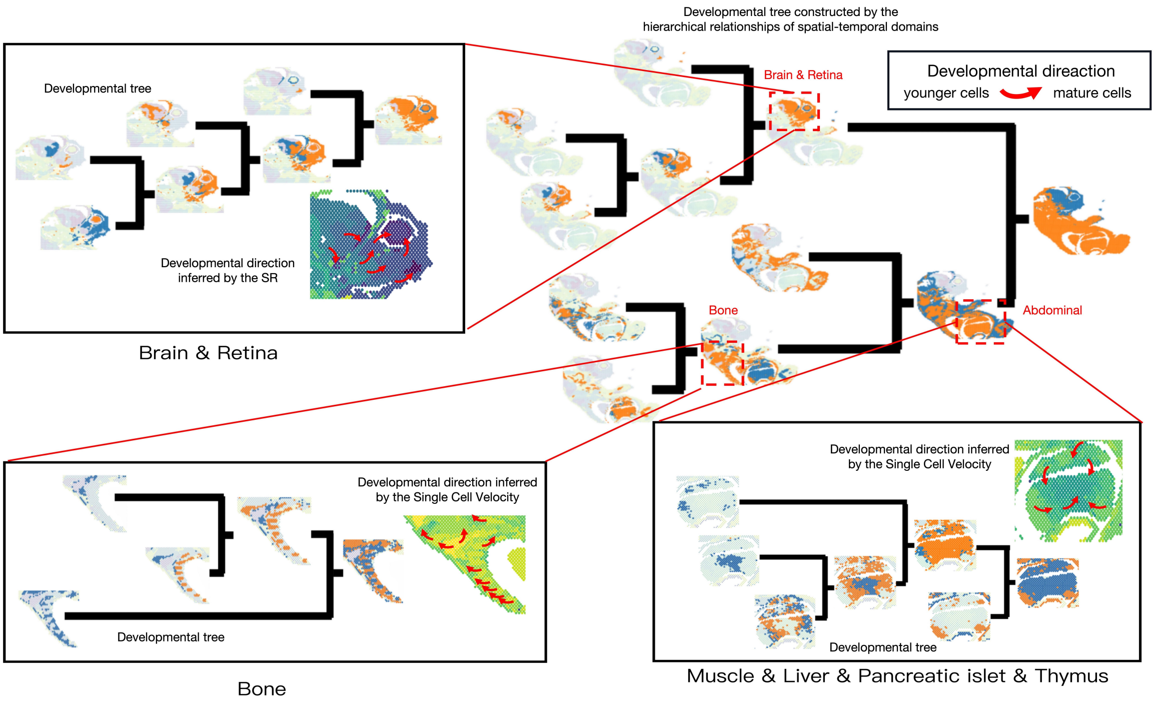


**Fig. S3. Hierarchical tree of domains visualizing the developmental path. Three regional traces are highlighted with average developmental levels and directions.**

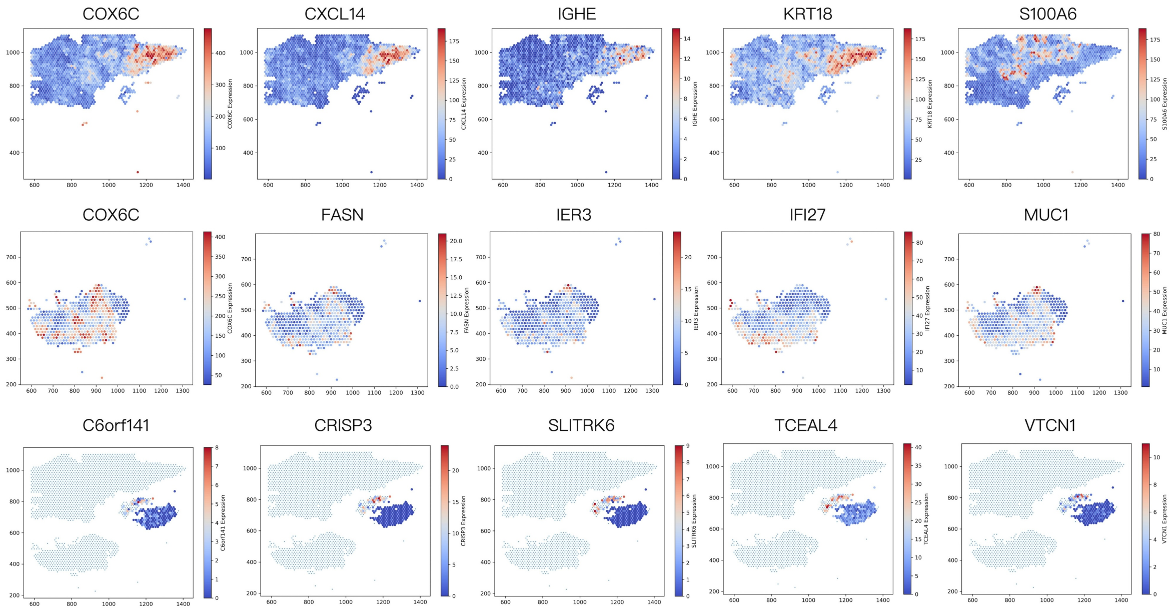


**Fig. S4. Spatial expression patterns of the top five differentially expressed genes identified in the infiltrating region. These genes were selected based on statistical significance and fold change across spatial-temporal domains. The visualization highlights their expression heterogeneity and potential functional relevance to cancer invasion dynamics along the developmental timeline. Detailed gene lists for each zone are provided in Supplementary Tables S2–S4.**

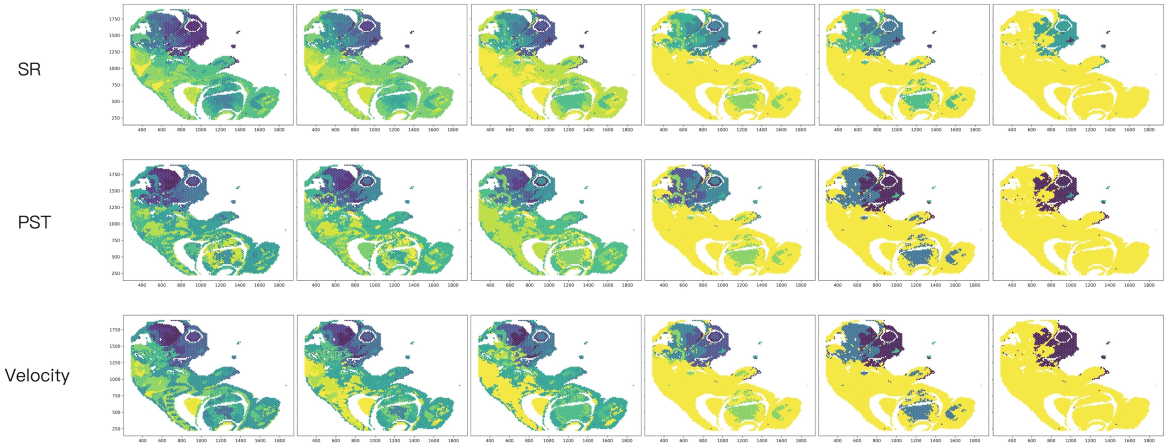


**Fig. S5. During the linkage process, the mean developmental level of each cluster was calculated based on the different cell maturity measurement methods: Signaling Entropy Rate, Pseudotime and Single Cell Velocity. The figure shows the distribution of average developmental levels across different Spatial-Temporal domains at six different stages in linkage process, with darker colors indicating more mature cells.**

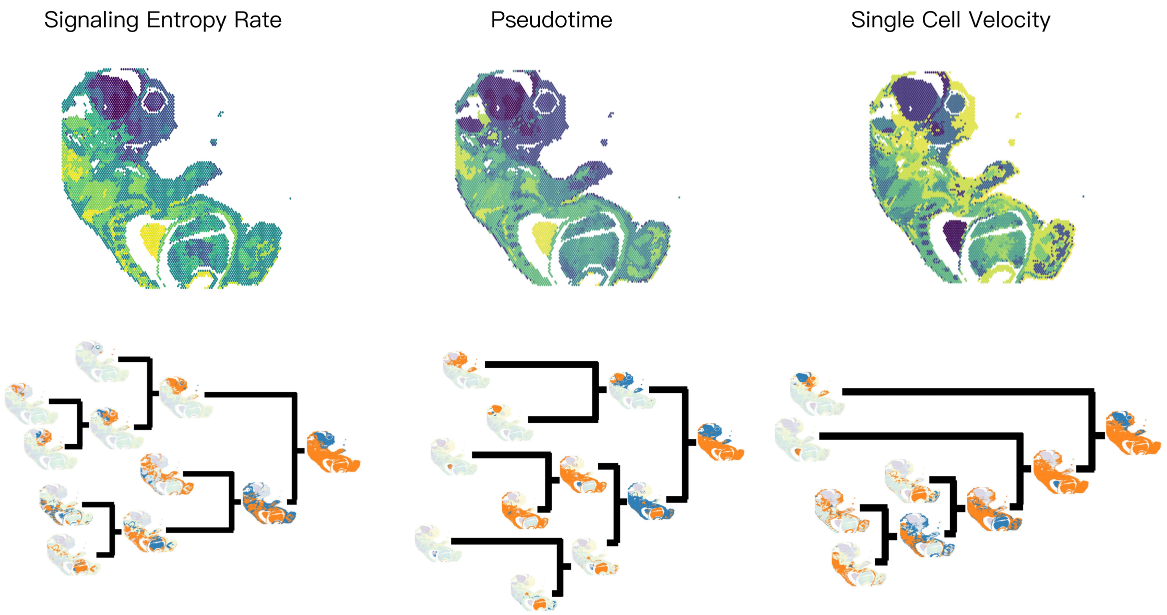


**Fig. S6. Comparison of stTrace results in mouse embryo based on different developmental level measurements: Signaling Entropy Rate, Pseudotime, and Single Cell Velocity. The figure includes spatial-temporal domain identification using stTrace and the developmental trees reconstructed from the hierarchical structure of the spatial-temporal domains.**

**Table 2.** The top 20 differentially expressed genes in infiltration region I.

| | baseMean | log2FoldChange | lfcSE | stat | pvalue | padj |
|---|---|---|---|---|---|---|
| **COX6C** | 56.11255326 | -0.050258069 | 0.002278257 | -22.05987333 | 7.68E-108 | 6.95E-104 |
| **CXCL14** | 3.63071774 | -0.073057108 | 0.003464232 | -21.08897709 | 1.00E-98 | 4.55E-95 |
| **S100A6** | 30.98703605 | 0.060038714 | 0.003854284 | 15.57713937 | 1.04E-54 | 3.14E-51 |
| **KRT18** | 52.9982942 | -0.020097039 | 0.001411588 | -14.23718614 | 5.39E-46 | 1.22E-42 |
| **IGHE** | 1.97783047 | -0.060196248 | 0.004322694 | -13.92563116 | 4.43E-44 | 8.02E-41 |
| **VSTM2A** | 0.943687705 | -0.074974549 | 0.005738393 | -13.06542649 | 5.19E-39 | 7.83E-36 |
| **HSP90AB1** | 15.8484957 | -0.020759048 | 0.00169357 | -12.25756555 | 1.53E-34 | 1.98E-31 |
| **PRDX1** | 6.483607034 | -0.025912254 | 0.002134967 | -12.13707504 | 6.72E-34 | 7.61E-31 |
| **SLC39A6** | 3.376240533 | -0.038285661 | 0.003258866 | -11.74815513 | 7.22E-32 | 7.26E-29 |
| **FASN** | 7.561142784 | -0.029598098 | 0.00256503 | -11.53908667 | 8.38E-31 | 7.59E-28 |
| **FN1** | 18.67297255 | 0.050478955 | 0.004474546 | 11.28135917 | 1.62E-29 | 1.34E-26 |
| **SCD** | 4.159107437 | -0.046923256 | 0.004210471 | -11.14441845 | 7.62E-29 | 5.75E-26 |
| **SLC7A2** | 1.478181974 | -0.051255044 | 0.0046223 | -11.08864532 | 1.42E-28 | 9.92E-26 |
| **PDCD4** | 1.058314097 | -0.054350639 | 0.004931137 | -11.0219282 | 3.00E-28 | 1.94E-25 |
| **LGALS1** | 10.50003869 | 0.033958753 | 0.003190365 | 10.64415757 | 1.86E-26 | 1.12E-23 |
| **AEBP1** | 6.664889756 | 0.051087087 | 0.004802196 | 10.63827663 | 1.98E-26 | 1.12E-23 |
| **PGK1** | 9.841811358 | 0.047083562 | 0.004496717 | 10.47065373 | 1.18E-25 | 6.28E-23 |
| **CA12** | 0.884787427 | -0.049974455 | 0.0047781 | -10.45906384 | 1.33E-25 | 6.70E-23 |
| **IFI27** | 47.33715637 | -0.021442607 | 0.002058538 | -10.41642628 | 2.09E-25 | 9.94E-23 |
| **SPP1** | 6.749024668 | 0.070879713 | 0.00691205 | 10.25451459 | 1.13E-24 | 5.11E-22 |

**Table 3.** The top 20 differentially expressed genes in infiltration region II.

| | baseMean | log2FoldChange | lfcSE | stat | pvalue | padj |
|---|---|---|---|---|---|---|
| **COX6C** | 125.5353711 | -0.059277153 | 0.006393815 | -9.271014302 | 1.84E-20 | 1.21E-16 |
| **MUC1** | 15.5127983 | -0.0606394 | 0.007883142 | -7.692288448 | 1.45E-14 | 4.73E-11 |
| **IFI27** | 19.45726099 | -0.056674287 | 0.007580545 | -7.476281231 | 7.65E-14 | 1.67E-10 |
| **IER3** | 4.60775369 | -0.080409191 | 0.010869062 | -7.397988248 | 1.38E-13 | 2.26E-10 |
| **FASN** | 5.090454335 | -0.069879333 | 0.010099544 | -6.919058546 | 4.55E-12 | 5.95E-09 |
| **MESP1** | 2.183505412 | -0.104285873 | 0.015643962 | -6.666205844 | 2.63E-11 | 2.86E-08 |
| **KRT19** | 53.02331568 | -0.037917257 | 0.005780173 | -6.559883042 | 5.38E-11 | 5.03E-08 |
| **TBC1D9** | 3.419888064 | -0.070422555 | 0.010830424 | -6.502289459 | 7.91E-11 | 6.47E-08 |
| **KRT18** | 21.43216745 | -0.039428894 | 0.006386101 | -6.174173395 | 6.65E-10 | 4.84E-07 |
| **SERF2** | 36.62758224 | -0.027474106 | 0.004552391 | -6.035094194 | 1.59E-09 | 1.04E-06 |
| **MGP** | 18.49715926 | -0.083878997 | 0.013993052 | -5.994331875 | 2.04E-09 | 1.21E-06 |
| **CLDN4** | 3.073535326 | -0.072002032 | 0.012038279 | -5.981090275 | 2.22E-09 | 1.21E-06 |
| **SRP14** | 8.23203652 | -0.045803199 | 0.007778141 | -5.888707582 | 3.89E-09 | 1.96E-06 |
| **YWHAZ** | 7.157341585 | -0.045413465 | 0.0077831 | -5.834881689 | 5.38E-09 | 2.33E-06 |
| **SLC9A3R1** | 7.366463008 | -0.048922733 | 0.00838969 | -5.831292205 | 5.50E-09 | 2.33E-06 |
| **SCD** | 1.847376402 | -0.097822037 | 0.016809627 | -5.819405486 | 5.91E-09 | 2.33E-06 |
| **UGCG** | 2.935338511 | -0.074776658 | 0.012859135 | -5.815061224 | 6.06E-09 | 2.33E-06 |
| **LY6E** | 4.687661887 | -0.056369079 | 0.009745737 | -5.783973028 | 7.30E-09 | 2.65E-06 |
| **MCCD1** | 1.6726446 | -0.088089948 | 0.015361104 | -5.734610425 | 9.77E-09 | 3.13E-06 |
| **S100A6** | 8.282430381 | -0.051622475 | 0.009007872 | -5.730817671 | 9.99E-09 | 3.13E-06 |

**Table 4.** The top 20 differentially expressed genes in infiltration region III.

| | baseMean | log2FoldChange | lfcSE | stat | pvalue | padj |
|---|---|---|---|---|---|---|
| **CRISP3** | 1.96968039 | -1.334059084 | 0.151868033 | -8.784331071 | 1.57E-18 | 6.99E-15 |
| **SLITRK6** | 0.869758199 | -0.913527765 | 0.143578692 | -6.362558061 | 1.98E-10 | 4.41E-07 |
| **TCEAL4** | 6.450498374 | -0.206155156 0.035776339 | -5.762332374 | 8.30E-09 | 1.23E-05 | |
| **VTCN1** | 0.771281401 | -0.446825697 | 0.088010256 | -5.076973016 | 3.83E-07 | 0.000426159 |
| **C6orf141** | 0.765116351 | -0.317052889 | 0.064384678 | -4.924353129 | 8.46E-07 | 0.000551319 |
| **MUC1** | 10.84922577 | -0.113215034 | 0.023002761 | -4.921801942 | 8.58E-07 | 0.000551319 |
| **KRT19** | 138.5583333 | 0.063770899 | 0.012963217 | 4.919373004 | 8.68E-07 | 0.000551319 |
| **S100G** | 9.299042389 | 0.167477243 | 0.034615697 | 4.838187741 | 1.31E-06 | 0.000728026 |
| **S100A16** | 7.734038107 | -0.115836132 | 0.0246496 | -4.699310879 | 2.61E-06 | 0.001289252 |
| **DEGS2** | 2.616893915 | -0.182880101 | 0.03953805 | -4.625420342 | 3.74E-06 | 0.001661716 |
| **PLAAT4** | 5.278621163 | -0.099548951 | 0.022711154 | -4.383262571 | 1.17E-05 | 0.004620193 |
| **LAPTM4B** | 7.753048286 | 0.126275197 | 0.028901531 | 4.36915253 | 1.25E-05 | 0.004620193 |
| **S100A13** | 8.103337509 | -0.11147868 | 0.025910361 | -4.30247494 | 1.69E-05 | 0.005533738 |
| **S100A14** | 5.59517778 | -0.119412034 | 0.027885048 | -4.282296156 | 1.85E-05 | 0.005533738 |
| **ELF3** | 7.242658448 | -0.118871344 | 0.027772496 | -4.280182263 | 1.87E-05 | 0.005533738 |
| **TMEM132A** | 1.04923573 | 0.238918351 | 0.057953445 | 4.122591009 | 3.75E-05 | 0.010407815 |
| **YPEL3** | 2.736356475 | -0.115208803 | 0.028056808 | -4.106269002 | 4.02E-05 | 0.010513763 |
| **MGP** | 49.42685984 | 0.11845499 | 0.029470229 | 4.019479848 | 5.83E-05 | 0.014403472 |
| **RPS12** | 23.02227858 | -0.07312489 | 0.018299648 | -3.995972529 | 6.44E-05 | 0.015073043 |
| **ADORA2A** | 0.476970179 | -0.29553249 | 0.074755689 | -3.953311016 | 7.71E-05 | NA |
